# Dissociating refreshing and elaboration and their impacts on memory

**DOI:** 10.1101/423277

**Authors:** Lea M. Bartsch, Vanessa M. Loaiza, Lutz Jäncke, Klaus Oberauer, Jarrod A. Lewis-Peacock

## Abstract

Maintenance of information in working memory (WM) is assumed to rely on *refreshing* and *elaboration*, but clear mechanistic descriptions of these cognitive processes are lacking, and it is unclear whether they are simply two labels for the same process. This fMRI study investigated the extent to which refreshing, elaboration, and repeating of items in WM are distinct neural processes with dissociable behavioral outcomes in WM and long-term memory (LTM). Multivariate pattern analyses of fMRI data revealed differentiable neural signatures for these processes, which we also replicated in an independent sample of older adults. In some cases, the degree of neural separation within an individual predicted their memory performance. Elaboration improved LTM, but not WM, and this benefit increased as its neural signature became more distinct from repetition. Refreshing had no impact on LTM, but did improve WM, although the neural discrimination of this process was not predictive of the degree of improvement. These results demonstrate that refreshing and elaboration are separate processes that differently contribute to memory performance.

**Highlights:**

- Repeated reading, refreshing, and elaboration are differentiable in brain activation patterns in both young and older adults.
- Elaboration selectively improved long-term memory for young adults, and the size of the benefit was related to the neural separability of elaboration from other processes.
- Older adults implemented a sub-optimal form of elaboration, and this may be a factor contributing to age-related deficits in long-term memory.

**Ethics statement:** The study was approved by the ethical review board of the canton of Zurich (BASEC-No. 2017-00190) and all subjects gave informed written consent in accordance with the Declaration of Helsinki.

**Data and code availability statement:** All behavioral data and analysis scripts can be assessed on the Open Science Framework (osf.io/p2h8b/). The fMRI data that support the findings of this study are available on request from the corresponding author, LMB. The fMRI data are not publicly available due to restrictions of the *Swiss Ethics Committees on research involving humans* regarding data containing information that could compromise the privacy of research participants.

## Introduction

Working memory (WM) is a system for holding a limited amount of information available for processing (Baddeley, 1986), whereas episodic long-term memory (LTM) stores information permanently with presumably unlimited capacity (Tulving, 1972). WM and LTM are highly correlated constructs, and models of their relation suggest that how information is processed in WM strongly affects how well it is maintained in LTM (D’Esposito & Postle, 2015; Eriksson, Vogel, Lansner, Bergström, & Nyberg, 2015; Lewis-Peacock & Postle, 2008; Ranganath, 2006; Ranganath & Blumenfeld, 2005; Ranganath, Cohen, & Brozinsky, 2005; Crowder, 1982; Melton, 1963; Nairne, 1990, 2002; Cowan, 1995; Oberauer, 2002). Two control processes on information in WM have been argued to contribute to encoding in episodic LTM: *refreshing* and *elaboration* (Bartsch, Singmann, & Oberauer, 2018; Camos & Portrat, 2015; Craik & Tulving, 1975; Gallo, Meadow, Johnson, & Foster, 2008; Johnson, Reeder, Raye, & Mitchell, 2002; Loaiza & McCabe, 2013). Refreshing is understood as briefly thinking of a stimulus just after it is no longer physically present but while its representation is still active (Johnson et al., 2002). Elaboration refers to enriching the representation of the stimulus with knowledge about it, for instance by combining words into a meaningful sentence, or creating a mental image of a word’s meaning.

The aim of the present study is to investigate (1) whether refreshing and elaboration are neurally and behaviorally distinguishable processes, (2) how they affect WM and episodic LTM performance, and (3) to what extent age differences in these processes are responsible for memory deficits in older adults.

### Refreshing and elaboration: Behavioral impacts on WM and LTM

Refreshing has been argued to improve WM (see Camos et al., 2018 for a review), but the evidence for this claim is mixed (Souza & Oberauer, 2017; Souza, Vergauwe, & Oberauer, 2018; Bartsch et al., 2018). Further, Refreshing has been claimed to also improve episodic LTM (Johnson et al., 2002), nevertheless, evidence for this is also ambiguous (Bartsch et al., 2018). Currently, there are two experimental approaches to investigate the effects of refreshing on memory: (a) Instructions to “think of” an item in WM, and (b) variations of free time during a WM task that could be used for refreshing. Both of these approaches are subject to an alternative interpretation: When instructed to “think of” a word, people are likely to think of its meaning, and this could include forming a mental image, and/or relating it to other WM contents – that is, they could elaborate the word. Likewise, when given free time, people could use it to elaborate rather than refresh the WM contents.

Elaboration has been shown to reliably improve episodic LTM (e.g. Craik & Tulving, 1975, Gallo, Meadow, Johnson, & Foster, 2008), whereas evidence for a benefit of elaboration for WM is more mixed: Correlational studies show a positive relationship between elaborative strategies and verbal WM recall (Bailey, Dunlosky, & Kane, 2008, 2011; Dunlosky & Kane, 2007; Kaakinen & Hyönä, 2007), and some experimental work has shown that semantic compared to shallow processing of the memoranda yields greater WM recall (Loaiza, McCabe, Youngblood, Rose, & Myerson, 2011; Rose, Buchsbaum, & Craik, 2014; Rose, Craik, & Buchsbaum, 2014). Conversely, other work has shown benefits of semantic processing only for episodic LTM but not WM (Loaiza & Camos, 2016; Rose & Craik, 2012; Rose et al., 2010). Bartsch, Singmann, and Oberauer (2018) and colleagues showed that elaboration benefited LTM, but refreshing did not, and neither elaboration nor refreshing benefited WM.

### Refreshing vs. elaboration: Neural correlates

Table 2 shows an overview of the reported brain regions associated with refreshing and/or elaboration. Refreshing has been associated with activity in the left dorsolateral prefrontal cortex (dlPFC, BA 8/9), and activity in the dlPFC during refreshing predicted subsequent LTM for the refreshed information (Johnson, Raye, Mitchell, Greene, & Adam, 2003; Raye, Johnson, Mitchell, Greene, & Johnson, 2007; Raye, Johnson, Mitchell, Reeder, & Greene, 2002). A meta-analysis (Johnson et al., 2005) identified frontal regions, specifically left dlPFC (BA 9/46), ventrolateral PFC (vlPFC, BA 44/45/47), and the left anterior PFC (BA 10) as associates of refreshing various stimulus materials.

Although the dlPFC has been suggested to underlie refreshing, its activation has also been shown to predict subsequent LTM in studies of elaboration (or “relational encoding”) wherein the semantic relationship between two items is elaborated upon (e.g. Blumenfeld & Ranganath, 2007). For the ease of the reader, we will refer to *relational encoding* as *elaboration* from now on. The neural correlates of elaboration have not always been specific or limited to the dlPFC: earlier studies have more generally associated the lateral PFC with semantic elaboration (e.g., Kapur et al., 1994; Wagner et al., 1998) and relational elaboration (e.g., Addis & McAndrews, 2006; Fletcher, Shallice, & Dolan, 2000; Murray & Ranganath, 2007). Yet, numerous studies have associated the dlPFC with elaboration, showing higher activation in the dlPFC when people elaborate, and that this activation also predicts subsequent memory (Blumenfeld, Parks, & Yonelinas, 2010; Blumenfeld & Ranganath, 2007; Davachi, Maril, & Wagner, 2001; Ragland et al., 2012). Collectively, this evidence suggests that elaboration of the memoranda in WM is what makes the dlPFC important for LTM.

Despite the neural similarities observed for refreshing and elaboration, there are also differences, which could be due to genuine differences between these processes, or to dissimilarities in the methods used to study these processes. First, the neural correlates of refreshing have been studied for single items only, with no instructed elaboration (e.g. Johnson et al., 2005; Raye et al., 2007, 2002), and this item-specific neural processing was localized almost exclusively to left lateral dlPFC. Conversely, elaboration studies have used multiple items, such as pairs (Blumenfeld et al., 2010) or triplets of words (e.g. Blumenfeld, 2006; Davachi, Maril, & Wagner, 2001), and localized the associated activity to the bilateral dlPFC. Second, the refreshing studies have relied on incidental encoding, wherein participants are not informed of the upcoming memory test, whereas the elaboration studies employ intentional encoding. Therefore, clarifying the underlying neural processes of refreshing and elaboration requires greater consistency between the methods used to investigate them.

### Refreshing and elaboration: Age effects

Past research has provided extensive evidence that episodic LTM declines with age (e.g., Hoyer & Verhaeghen, 2006; Naveh-Benjamin & Old, 2008; Zacks, Hasher, & Li, 2000), but the source of the deficit is still under debate. One view is that WM maintenance processes and recruitment of corresponding brain areas decline in older age (Hoareau, Lemaire, Portrat, & Plancher, 2016; Plancher, Boyer, Lemaire, & Portrat, 2017; Smith, 1980). For instance, it has been shown that older adults exhibit reduced refreshing-related brain activity in the left dlPFC and reduced refreshing benefits for episodic LTM relative to young adults (Johnson, Mitchell, Raye, & Greene, 2004; Raye, Mitchell, Reeder, Greene, & Johnson, 2008). Another possibility is that older adults are less likely than younger adults to engage in elaboration, thereby resulting in deficient retention (Smith, 1980). For example, some work has shown older adults are able to capitalize on experiment-administered elaborative strategies but show deficiencies in generating elaborative strategies themselves (Rankin & Collins, 1985, see also Kamp & Zimmer, 2015). A meta-analysis reported that age-related differences in subsequent memory are associated with under-recruitment of the occipital and fusiform cortex in older adults, as well as an over-recruitment of medial and lateral regions of PFC and parietal lobe (Maillet & Rajah, 2014). These findings suggest inefficient recruitment of brain regions that are important for elaboration, thereby leading to age-related memory deficits.

### The present study

The goal of the present study was to investigate to what extent elaboration and refreshing are separable processes, given their neural overlap as well as their similar proposed beneficial effects for memory. So far, only one study has investigated both processes in one experiment, and the behavioral results demonstrated that the processes have divergent contributions to LTM (Bartsch et al., 2018). We aimed at extending this previous study by not only investigating whether refreshing and elaboration are distinct in their contribution to WM and LTM formation, but also whether they are supported by separable neural activation patterns. Furthermore, we aimed to investigate their contribution to age-related memory deficits.

We applied multivariate pattern analyses (MVPA; e.g., Haxby, Connolly, & Guntupalli, 2014; Haxby et al., 2001; Haynes & Rees, 2006; Lewis-Peacock & Norman, 2014; Norman, Polyn, Detre, & Haxby, 2006) to fMRI data of young adults and older adults performing the word list encoding task of Bartsch et al. (2018). This analysis approach allowed us to evaluate whether brain activity patterns associated with refreshing items and with elaborating items in WM could be differentiated. These neural measures were then linked to behavioral outcomes on tests of both WM and LTM. MVPA is especially sensitive to detecting fine-grained differences between neural activation patterns that are not detectable using conventional analyses (Kriegeskorte & Bandettini, 2007).

If refreshing and elaboration are two labels for the same process, then the pattern of behavioral effects should be similar for WM and on LTM, and the patterns of brain activity supporting these processes should be indistinguishable. If refreshing and elaboration are distinct processes, they should have different behavioral effects and separable patterns of neural activation.

## Method

### Subjects and general procedure

We recruited 30 healthy, right-handed young adults (15 females; mean age = 24.2, SD = 2.97 years) from the student population of the University of Zurich as well as 27 healthy, right-handed older adults from the community (13 females; mean age = 69, SD = 3.47 years). None of the participants had taken part in our prior study (Bartsch et al., 2018). Handedness was measured through observation of the writing hand. Subjects were screened for their ability to undergo a magnetic resonance imaging session. Furthermore, they completed the Digit–Symbol Substitution test (DSS; Wechsler, 1982), serving as an indicator of processing speed, and the mini-mental-status examination (MMSE; Folstein, Folstein, & McHugh, 1975) to screen for cognitive impairment. We ensured that all subjects’ MMSE score was above 25. All subjects performed a WM task while being scanned with a 3-T MRI scanner, and subsequently an LTM task outside the scanner. The session ended with a computerized version of a vocabulary test (Lehrl, 2005), a marker test for crystallized intelligence. Table 1 shows the descriptive statistics of the age differences in our sample, indicating that our sample of young and old adults show typical differences in measures of crystallized intelligence and processing speed (Li et al., 2004). The study was approved by the ethical review board of the canton of Zurich and all subjects gave informed written consent in accordance with the Declaration of Helsinki. The participants were compensated with either 60 Swiss Francs (about 60 USD) or partial course credit for the two-hour session.

**Table 1.**
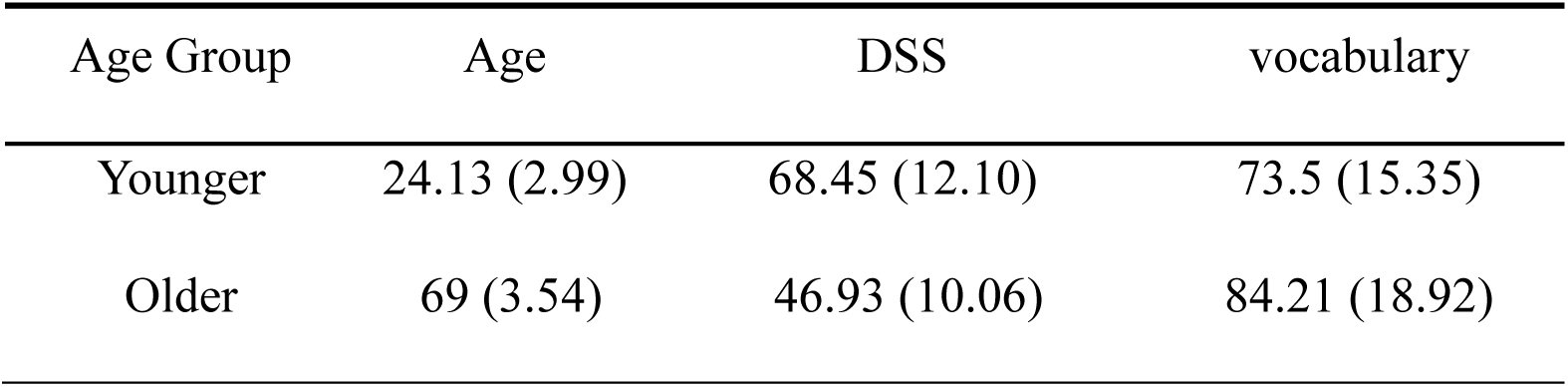
Sample Description (means (and standard deviations))

**Table 2.** ROIs with their corresponding BAs and references of previous reporting in univariate analyses in the literature.

| Label | sub region | BA | Labeled region reported in |
| --- | --- | --- | --- |
| frontal | inferior frontal | 44,45,47 | Johnson et al., 2005;<br>Johnson, Mitchell, Raye, & Greene, 2004;<br>Raye, Johnson, Mitchell, Greene, &<br>Johnson, 2007; |
|  | middle &<br>superior frontal | 4,6,<br>8,9,10,46 | Raye, Mitchell, Reeder, Greene, &<br>Johnson, 2008<br>Blumenfeld, 2006;<br>Blumenfeld, Parks, & Yonelinas, 2010;<br>Kim & Giovanello, 2011; Murray &<br>Ranganath, 2007 |
| parietal |  | 3,7,40 | Johnson et al., 2004;<br>Kim & Giovanello, 2011;<br>Murray & Ranganath, 2007;<br>Raye, Johnson, Mitchell, Reeder, &<br>Greene, 2002;<br>Raye et al., 2007, 2008 |
| fusiform |  | 19, 37 | Murray & Ranganath, 2007;<br>Raye et al., 2008 |

### Paradigm

The paradigm is the same as reported in a recent study (Bartsch et al., 2018), adapted for use in the MRI scanner. We asked participants to remember six nouns in serial order (see Figure 1). After list presentation, either the first three words or the last three words were to be processed again in one of four ways, depending on the experimental condition. During encoding it was not predictable which half of the items would have to be processed. In the *repeat* condition, the three words appeared again sequentially on the screen, and the subjects had to simply re-read them silently. In the *refreshing* condition, the to-be-processed words were replaced by refreshing prompts appearing at the same location. The subjects were instructed to “think of” the corresponding words as soon as the prompts were shown. In the *elaboration* condition, the three to-be-processed words were shown again sequentially on the screen, and subjects were instructed to generate a vivid mental image of the three objects interacting. The stimuli appearing on the screen in that condition did not differ from the repeat condition, leaving the instruction to form a vivid mental image as the only difference between these conditions. Finally, in the combined *refreshing with elaboration* condition, the participants had to “think of” the words replaced by the prompts, and additionally form a vivid mental image of those items. Again, the event sequence of this condition does not differ from the refreshing condition apart from the instruction to form a mental image. Memory was tested with a four-alternatives forced-choice task, which we describe in detail below (see section Procedure: Working memory task).

**Figure 1.**
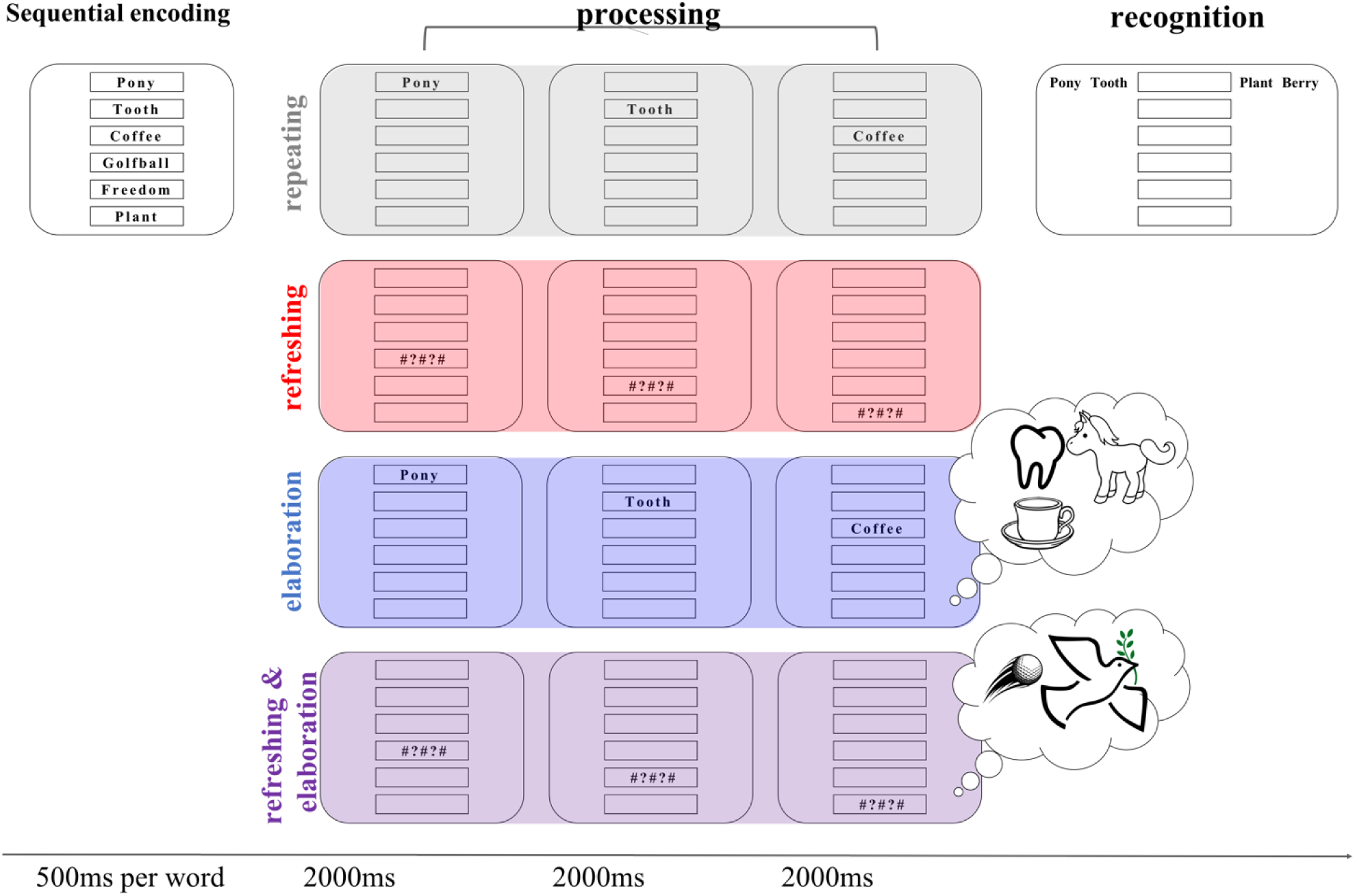
Illustration of the immediate memory paradigm. Subjects were shown a list of six words sequentially, followed by either the first or second triplet being processed according to the four experimental conditions. The trial ended with a recognition test in which each list item was tested in their order of presentation using a 4-alternative forced-choice procedure. The grey panel shows the repeating condition with an example of the first triplet being processed. The red panel shows the refreshing condition, here for the second triplet to be processed. The blue and purple panels show the elaboration and refreshing with elaboration condition, respectively, which were preceded by the instruction to form a mental image, but which were equivalent in the visual input to the respective conditions without the elaboration instruction.

The experiment used a 2 × 2 × 2 × 2 (repeat/refresh [repeat, refresh] × elaboration [with elaboration, without elaboration] × processing [processed triplet, unprocessed triplet] × age [young adults, older adults]) within-subject, between-age group design. As in the studies of Raye and colleagues, we compared instructed refreshing to a repeating (re-reading) baseline during the maintenance phase of a WM task (e.g. Raye et al., 2002). In the two additional conditions we instructed participants to elaborate a subset of the items they held in memory. Elaboration logically entails attending to the words, either in memory or in the environment. When elaboration is applied to words just encoded into WM, but no longer presented, it entails refreshing, whereas when elaboration is applied to words while they are presented, it entails (re-)reading, as in the *repeat* condition. Therefore, we implemented two elaboration conditions: One in which words were repeated and elaborated, and one in which they were refreshed and elaborated. Contrasting these two conditions allowed us to gauge any unique effects of elaboration. In addition, this design allowed us to evaluate whether combining elaboration with refreshing is more effective than either of them alone.

How can we measure the effect of refreshing in our paradigm? The Johnson et al. studies – testing the effect of refreshing on LTM – used *repeat* as the baseline, and therefore we followed their precedent for assessing the effect of refreshing on LTM. For assessing the effect of refreshing on WM, the *repeat* condition is not a suitable baseline because it provided a second chance for encoding the word into WM. Therefore, we assessed the effect of refreshing against the baseline used in Souza, Rerko, & Oberauer, 2015 (i.e., comparison within the memory set between items refreshed more vs. less) by comparing the items that were processed in refreshing trials to the items within the same trial that were not processed further after initial encoding.

#### Materials

The stimuli were nouns randomly drawn from a pool of 863 German abstract and concrete nouns for each subject. The nouns were between three and 15 letters long and had a mean normalized lemma frequency of 30.81/million (drawn from the dlexdb.de lexical database).

#### Procedure

##### Working memory task

The sequence of an experimental trial is illustrated in Figure 1. The experiment was performed using Presentation® software (Version 18.0, Neurobehavioral Systems, Inc., Berkeley, CA, www.neurobs.com). The six to-be-remembered words in each trial were sequentially presented in boxes from top to bottom on the screen, each for 500 ms. A processing cue was presented 1000 ms after the last memory item, indicating whether the first half or the second half of the list had to be processed again, and in which way. In the repeat and elaboration conditions, each word in the to-be-processed triplet was shown again for 1400 ms, followed by a 600 ms inter-stimulus interval. In the refreshing and refreshing-with-elaboration conditions, each to-be-processed word of a triplet was replaced by a refreshing prompt (*#?#?#*) in its corresponding box, and participants were instructed to “think of” the word in that box. In the elaboration and refreshing-with-elaboration conditions, participants were additionally instructed to form a vivid mental image of the three words interacting with each other.^1^

After processing the words in the cued triplet, participants’ memory for each list item was tested in their order of presentation using a 4-alternative forced-choice procedure. For each tested item, four words were presented from which the subject could choose the correct word in the currently tested list position with a button press. All test sets included the following four response options: the target (i.e., correct) word, one lure from the same triplet of words within the present list, one lure from the other triplet of the present list, and one new word. This choice had to be made for each of the serial positions successively and with a time limit of 2500 ms for the young and 3500 ms for the older adults per serial position to ensure controlled timing for the fMRI image acquisition. We applied this 4-alternative forced-choice recognition task in order to test both memory for items (i.e., discriminating between items that have been presented in the current memory list and new items) and for serial order (i.e., discriminating between the item in the tested position and other list items). A variable 8 – 10 second inter-trial interval preceded the start of each trial.

Within each block of four trials, the same type of processing was instructed throughout, and a screen repeating the instructions of the particular condition was shown prior to the beginning of each block. The order of the condition blocks was randomized between subjects. Each of the four fMRI runs consisted of four blocks, one for each condition (with 4 trials per condition (repeat, refreshing, elaboration, and refreshing with elaboration), and 4 trials per block as described above). In total, each subject performed 64 trials across 4 runs, with 16 trials per condition (repeat, refreshing, elaboration, and refreshing with elaboration), and 4 trials per condition per run.

##### Long-term memory task

After leaving the scanner participants were brought into a separate room, where they performed the computerized LTM task. We assessed participants’ LTM for the words they had encoded for the WM tests throughout the experiment. To this end, we presented in each trial the first word of a triplet from one of the studied memory lists. We asked participants to choose, from four different options, the word that had followed the target word in that triplet. The probe words included the correct word (i.e., which could be either the word in the second or third position of the target triplet for the first prompt, and the fifth or sixth word for the second prompt), two words from another list, and a new word. This allowed us to keep the format of the LTM test very similar to the WM test, and furthermore to compare in each trial the memory performance for words from the processed versus unprocessed triplets. As in the WM test, the LTM test also provided information about both item memory (i.e., which words have been presented in the experiment) and relational memory (i.e., which words have been together in a triplet). The participants were made aware of the LTM test before the start of the experiment.

### fMRI Data Acquisition and Preprocessing

Whole brain images were acquired with the 3 T Philips Ingenia MRI scanner with a 32-channel head coil, located at the University Hospital Zurich, Switzerland. High-resolution T1-weighted images were acquired for all subjects with a Turbo field echo (TFE) sequence (8ms time repetition (TR), 3.7ms time echo (TE), 8° flip angle, 160 sagittal slices, 240 × 240 inplane, mm isotropic). Blood oxygen level-dependent (BOLD)-sensitive functional MRI data were acquired using a gradient-echo, echo planar sequence (2 s TR, 35ms TE) within a 72 × 70 matrix (32 transverse slices, 3 mm isotropic).

Following the acquisition of the structural images, four MRI acquisition runs were collected for each subject, in which they performed a 10-min block of a six-item WM task with a processing delay. fMRI data preprocessing (slice-time correction and realignment) was performed with SPM12 (Penny, Friston, Ashburner, Kiebel, & Nichols, 2011). Subjects’ functional scans were aligned by realigning the first volume in each run to the first volume of the first run, and then registering each image in each run to the first volume of that run. The middle functional slice served as a reference for slice-time correction. Further, the functional volumes were co-registered to the T1 anatomical image. No spatial smoothing was imposed on the data and the data were analyzed in each subject’s native space, since MVPA is known to capture fine-grained patterns in the brain activations, which would be blurred by spatial smoothing (e.g. Brauchli, Leipold, & Jäncke, 2019).

### Analysis of Behavioral Data

All data and analysis scripts can be assessed on the Open Science Framework (osf.io/p2h8b/). We analyzed the behavioral data using a Bayesian generalized linear mixed model (BGLMM) implemented in the R package *rstanarm* (Stan Development Team, 2018) following the exact analysis pipeline reported by Bartsch and colleagues (2018).^2^ The dependent variable was the number of correct and incorrect responses in each cell of the design per participant. Correct responses were defined as choosing the target item from the four alternatives. Therefore, we assumed a binomial data distribution predicted by a linear model through a probit link function (i.e., a mixed effects probit regression). The fixed effects were processing (processed versus not-processed triplet), repeat/refresh (repeated versus refreshed items), elaboration (with versus without elaboration instruction), age (young vs. older adults), and all their interactions. Following the recommendation of Barr and colleagues (Barr, Levy, Scheepers, & Tily, 2013; see also Schielzeth & Forstmeier, 2009), we implemented the maximal random-effects structure justified by the design; by-participant random intercepts and by-participant random slopes for the fixed effects excluding age (as these factors were within-subject factors). In addition, we estimated the correlation among the random-effects parameters. As factor coding, we used the orthonormal contrasts described in Rouder, Morey, Speckman, and Province (2012); section 7.2) that guarantee that priors affect all factor levels equally. For factors with two levels as employed here this corresponds to contrasts with values of √2/2 and −√2/2. Following Gelman et al. (2013), the regression coefficients were given weakly informative Cauchy priors with location 0 and scale 5. We used completely non-informative priors for the correlation matrices, so-called LKJ priors with shape parameter 1 (Stan Development Team, 2018).

Bayesian procedures provide posterior probability distributions of the model parameters (i.e., the regression weights) that express uncertainty about the estimated parameters. The highest density regions (HDRs) of these posteriors can be used for statistical inference. A 95% HDR represents the range in which the true value of a parameter lies with probability 0.95, given model and data (Morey, Hoekstra, Rouder, Lee, & Wagenmakers, 2016). If zero lies outside the Bayesian HDR there is strong evidence for the existence of the corresponding effect. Although the strength of evidence varies continuously, for simplicity we will describe effects as “credible” if their HDRs exclude zero. We used an MCMC algorithm (implemented in Stan; Carpenter et al., 2017) that estimated the posteriors by sampling parameter values proportional to the product of prior and likelihood. These samples are generated through 4 independent Markov chains, with 1000 warmup samples each, followed by 1000 samples drawn from the posterior distribution which were retained for analysis. Following Gelman and colleagues (2013), we confirmed that the 4 chains converged to the same posterior distribution by verifying that the 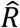 statistic – reflecting the ratio of between-chain variance to within-chain variance – was < 1.01 for all parameters, and we visually inspected the chains for convergence.

### Generation of ROIs

We included voxels within a distributed mask of ROIs – encompassing frontal, fusiform and parietal regions – that were previously reported in fMRI studies investigating either refreshing or elaboration and that had shown subsequent memory effects and/or significant activation differences between repeating and refreshing or elaboration in univariate analyses (see Table 1 for details). The search for those ROIs was performed using the neurosynth.org database and keyword-based search in pubmed.gov. Anatomical ROIs were generated using an automated parcellation method from *FreeSurfer*. Briefly, a surface mesh model was reconstructed for each subject’s brain. Each subject’s surface was then auto-parcellated based on the folding pattern of the gyri and sulci. We constructed the combined *frontal-fusiform-parietal* mask using *fslmaths*, encompassing Brodmann areas 4, 6, 8, 9, 10, 44, 45, 46 and 47 for the frontal mask, and Brodmann area 3, 7 and 40 for the parietal mask. The fusiform mask consisted of the fusiform label of the *aparc* atlas.

### Multivariate Pattern Analyses of fMRI Data

MVPA provides greater inferential power than classical univariate approaches due to its higher sensitivity at detecting information in neural signals. As a result, MVPA has led to the successful within-category decoding of the contents of WM at an item level (LaRocque, Riggall, Emrich, & Postle, 2017) as well as the characterization of neural representations in different states of WM (Christophel, Iamshchinina, Yan, Allefeld, & Haynes, 2018; Lewis-Peacock, Drysdale, Oberauer, & Postle, 2012; Lewis-Peacock, Drysdale, & Postle, 2015; Rose et al., 2016). The sensitivity of MVPA was further established by a study demonstrating that allegedly category-selective brain regions detected in univariate analyses of the BOLD signal during the delay period of a WM task still carried patterns of activity associated with another category of information that was currently relevant for behavior (Lewis-Peacock & Postle, 2012).

MVPA was performed in MATLAB using the Princeton MVPA toolbox (http://code.google.com/p/princeton-mvpa-toolbox). The classification algorithm used for this analysis was a L2-regularized binary logistic regression (predicting one category vs. the others), that uses Carl Rasmussen’s conjugate gradient minimization algorithm, with a penalty term of 50. The classification was performed in the anatomically defined ROI (the frontal-fusiform-parietal mask) defined above. All neural data were high-pass filtered with a cut-off of 128 seconds and z-scored across trials, within runs, before running MVPA. We performed ANOVA-based feature selection of all active voxels within the ROI and chose the voxels that individually were able to discriminate between the four conditions (repeat, refreshing, elaboration and refreshing with elaboration) significantly (*p* < .05) over the course of the experiment. This univariate feature selection technique has been shown to reliably improve classification accuracy in MVPA of fMRI (Lewis-Peacock & Norman, 2014). The ANOVA feature selection is a mass-univariate method that looks for voxels with significant differences in mean value across conditions (e.g. A is different from B C D). This is done using an ANOVA to test the null hypothesis at each voxel that the mean voxel value for the experimental conditions is the same. The score is the p-value which is used for thresholding (e.g., selecting voxels with p < .05; for further explanation, see Pereira, Mitchell, & Botvinick, 2009). To avoid circularity in the data analysis, feature selection was performed separately for each iteration of the cross-validation classifier training algorithm, using independent training and testing sets in each iteration (Kriegeskorte, Simmons, Bellgowan, & Baker, 2009). For each fold of the cross-validation analysis within each subject, the classifier was trained on 3 runs (48 trials, 12 per condition) and tested on 1 run (16 trials, 4 per condition).

For subjects who did not show successful classification in the larger ROI, we investigated whether any of the three subregions could differentiate the processes (i.e. frontal, fusiform and parietal). If this were the case, it would indicate that at least one of the subregions contributed more noise than signal to the classification problem. The number of participants for which subregion selection was used is reported in the Results. The average number of feature-selected voxels for young participants was 785.98 (SEM = 90.96) and 1095.3 (SEM = 172.11) for older participants. The pattern of activity across these voxels was used as the input to each participant’s pattern classifiers.

#### MVPA — multiple process discrimination

The classification procedure used k-fold cross-validation on the data from the WM task. Preprocessed fMRI data from each 6-s processing period (three volumes) from the WM trials, were used for the analysis. As is standard practice in MVPA (Lewis-Peacock & Norman, 2014), all trial regressors were shifted forward in time by 6 s to account for haemodynamic lag of the blood-oxygen-level dependent signal (typically estimated as 4–8 s to peak after event onset). Our analysis scheme incorporated each functional volume (acquired over a 2-s TR) as a separate training event, so that every trial resulted in three events corresponding to the three prompts to repeat, refresh or elaborate the memoranda. Each event was associated with an array of features corresponding to BOLD signals in voxels in the ROI being used.

We derived those three training samples from the consecutive TRs in the processing phase of each trial, rather than computing a voxel-wise average of three volumes from each trial, because there were three unique processing cues presented on each trial (e.g., [Pony] … [Tooth] … [Coffee]; see Figure 1). Although these measurements would be temporally correlated due to the sluggishness of the fMRI signal, there could be different levels of process engagement for each of these cues, reflected in different voxel-wise activity patterns at each TR, and thus we wanted the training data to capture these differences. For instance, elaboration by forming an integrative image is more likely to happen after cues to the second and third word to be elaborated, whereas refreshing can proceed on each word individually, so the two processes might differ in how they evolve over the three successive cues. Critically, using multiple (correlated) samples from each trial does not unfairly bias the cross-validation analysis which was used to assess classification accuracy. This is because the training set and testing set were completely independent. Training on correlated samples could lead to overfitting of the training data, if anything, which would reduce generalization to test data.

The k-fold cross-validation scheme (k = 4, for each of the runs) trained a classifier, separately for each participant, on the data of the four conditions (repeat, refreshing, elaboration and refreshing with elaboration) from three runs and then used this classifier to test the data from the withheld run. This process was repeated until every run had been held out for testing. Following this logic, for each fold of the cross-validation within each subject, the classifier was trained on twelve trials and tested on four.

The statistical significance of classifier accuracy was evaluated by performing permutation tests on relabeled training data, in each cross-validation fold, and comparing the resulting distribution of classifier accuracies to the true (unshuffled labels) classifier accuracy with a one-sample, one-tailed t-test. This analysis scheme was performed for the whole ROI and for the single masks of frontal, fusiform and parietal regions. Finally, classification performance was also assessed using receiver operating characteristic (ROC) curves, which rank the classification outputs according to their probability estimates (from strongly favoring the target class to favoring one of the three non-target classes) and chart the relationship between the classifier’s *true positive rate* (probability of correctly labeling examples of the target class) and *false positive rate* (probability of incorrectly labeling examples of the non-target class as the target class) across a range of decision boundaries. The area under the curve (AUC) indexes the mean accuracy with which a randomly chosen trial of the target class could be assigned to their correct classes (0.5 = random performance; 1.0 = perfect performance).

#### MVPA — discrimination from repeat

In order to assess how the neural classification of the refreshing process as well as the neural classification of the elaboration process relates to an individual’s task performance, we used the repeat condition as a reference. First, we extracted classification scores from repeat and refresh trials only, using the classifiers that were trained on all four processes. The same was done for the perceptually identical conditions of repeat and elaboration. Once again, we assessed classifier performance for each binary classification problem using AUC. To evaluate whether the degree of neural separability between the conditions relates to the individuals’ memory performance, we performed a logistic regression relating the evidence values from the trained classifier for the respective condition on each trial to the WM outcome of that trial. To increase statistical power for this analysis, we performed a non-parametric bootstrap analysis using data sampled from all participants (see Lewis-Peacock, Cohen, & Norman, 2016).

#### Researcher Degrees of Freedom

Analyses of neural data involve many decisions, and when these decisions are informed by the data to be analyzed, there is a risk that they are biased in favor of a desired outcome (Simmons, Nelson, & Simonsohn, 2011). Some aspects of our analysis plan (in particular, the decision to use an anatomically defined ROI for the MVPA analyses, rather than whole-brain classification or searchlight analysis) were informed by the data of the young adults. Our analysis of the older adults’ data, however, used the exact same analysis pipeline as that for the young adults without any adjustment informed by the older adults’ data. Therefore, any convergent finding in both age groups can be thought of as having been directly replicated in a different population. For any finding that differs between age groups, there remains an ambiguity as to whether the divergence reflects a failure to replicate the finding in the young-adult sample, or a genuine age difference. Resolving this ambiguity requires a replication of the entire study with the present analysis plan.

## Results

### Behavioral Results

We replicated all effects of the young adults reported in a previous study (Bartsch et al. 2018). Figure 2 shows the estimated proportion of correct responses and their corresponding 95% highest posterior density regions for the immediate and delayed memory data. The posterior effect estimates are presented in Table 3 and Table 4. A first question was whether our manipulation of processing half of a memory list had an effect on memory. The credible main effect of processing on immediate and delayed memory supported an effect of our manipulation: Participants had better memory for items that were processed again after initial encoding than for items from the unprocessed triplets (see Table 3 & Table 4 and Figure 2). There was also a main effect of age, such that older adults showed worse memory performance on tests of both WM and LTM.

**Figure 2.**
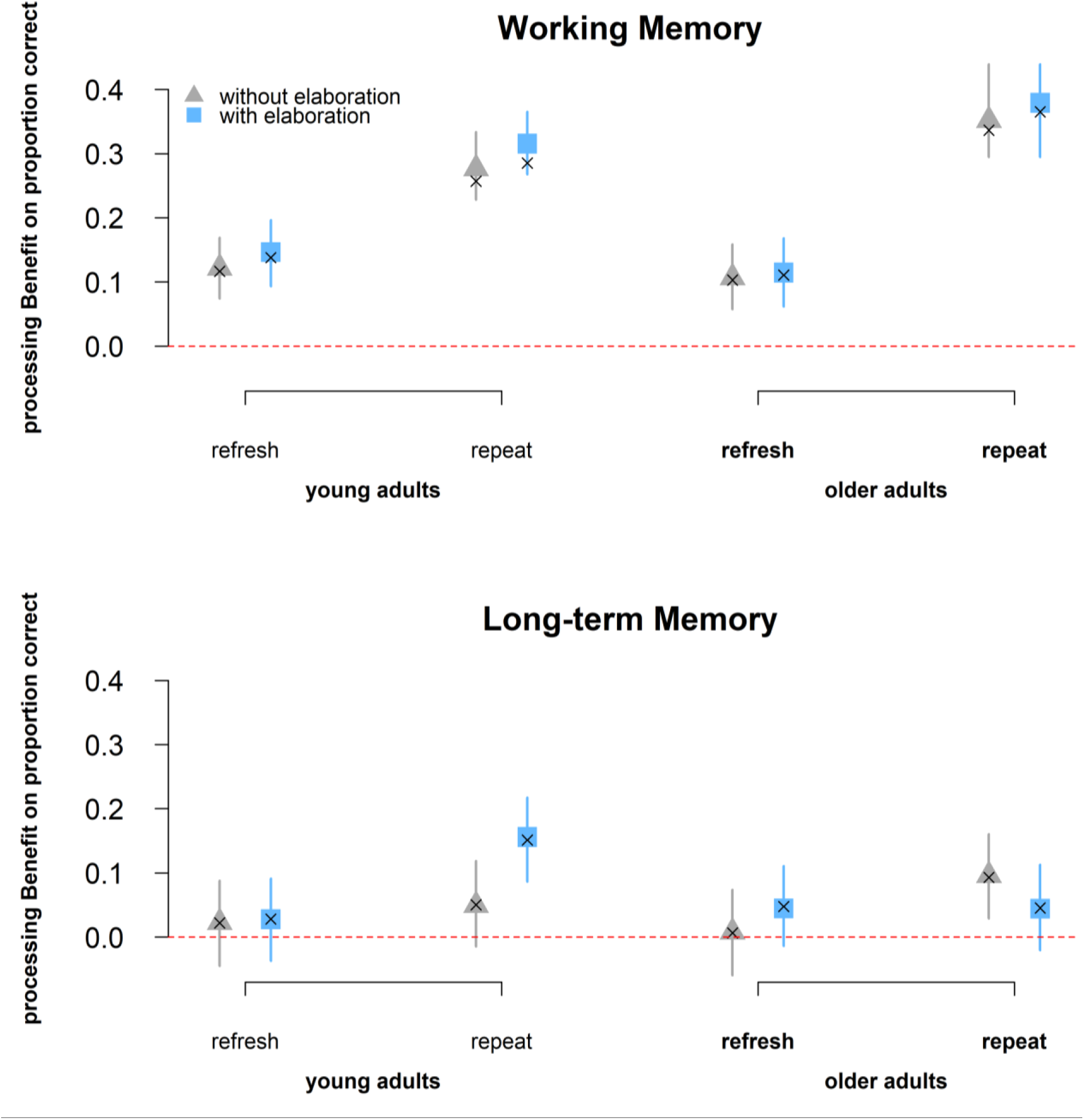
processing benefit in the WM (upper graph) and LTM (lower graph) task. The blue symbols and error bars represent estimated processing benefits and their 95% HDRs from the BGLMM for the conditions with elaboration, the grey symbols represent the same for the ones without elaboration. The crosses represent the observed means. Their overlap indicates that the model adequately describes the data. The red line represents the point of no difference in performance between the processed and the unprocessed triplet.

**Table 3.**
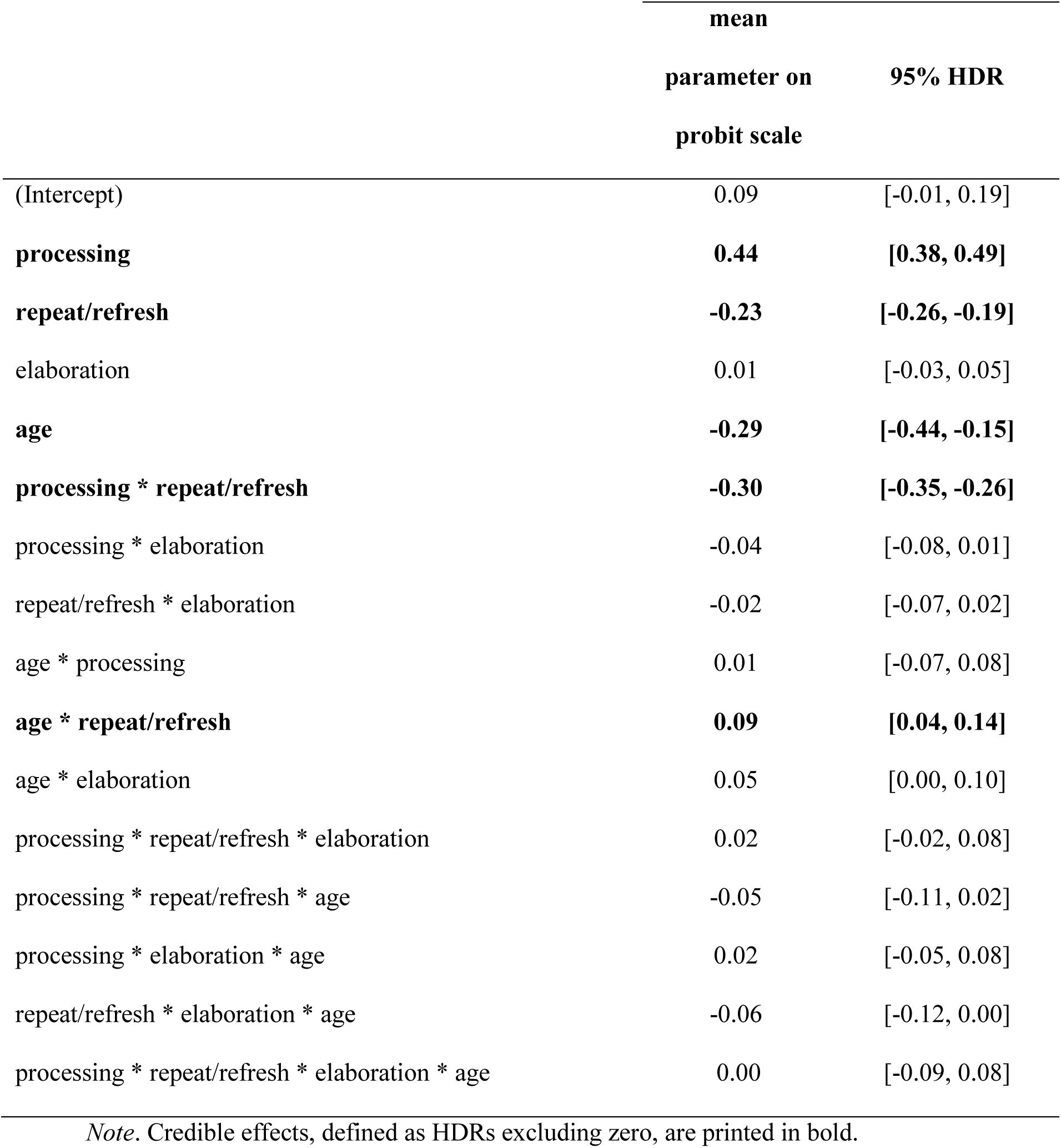
The posterior effect estimates and their 95% HDRs of the generalized linear mixed model for binomial response variables for the immediate serial memory data.

**Table 4.** The posterior effect estimates and their 95 % HDRs of the generalized linear mixed model for binomial response variables for the delayed memory data.

|  | mean |  |
| --- | --- | --- |
|  | parameter on | 95% HDR |
|  | probit scale |  |
| **(Intercept)** | **-0.29** | **[-0.38 -0.25]** |
| **processing** | **0.06** | **[0.01, 0.1]** |
| **repeat/refresh** | **-0.05** | **[-0.09, -0.01]** |
| elaboration | -0.04 | [-0.09, 0] |
| **age** | **-0.06** | **[-0.11, -0.01]** |
| processing * repeat/refresh | -0.04 | [-0.09, 0.02] |
| processing * elaboration | -0.01 | [-0.07, 0.05] |
| repeat/refresh * elaboration | 0.05 | [-0.01, 0.11] |
| age * processing | 0.03 | [-0.03, 0.09] |
| age * repeat/refresh | -0.05 | [-0.11, 0.01] |
| **age \* elaboration** | **0.08** | **[0.01, 0.14]** |
| processing * repeat/refresh * elaboration | -0.03 | [-0.12, 0.05] |
| processing * repeat/refresh * age | 0.05 | [-0.03, 0.14] |
| processing * elaboration * age | 0.05 | [-0.04, 0.13] |
| repeat/refresh * elaboration * age | -0.04 | [-0.13, 0.05] |
| processing * repeat/refresh * elaboration * age | -0.04 | [-0.15, 0.08] |

*Note.* Credible effects, defined as HDRs excluding zero, are printed in bold.

#### Working memory performance

We first tested how the effect of refreshing a subset of words in WM compares to the effect of repeated reading of these words. This is the comparison through which Johnson and colleagues evaluated the effect of refreshing on delayed memory (Johnson et al., 2002; Raye et al., 2007). There was a main effect of repeat/refresh (Table 3), but with an advantage of repeating over refreshing. This main effect was further qualified by the two-way interaction of processing and repeat/refresh, indicating that repeated words benefited more from being processed again than refreshed words did. Nevertheless, a beneficial effect of processing was found for both repeated words (Δ = 0.34, 95% HDR = [0.31, 0.37]) and refreshed words (Δ = 0.12, 95% HDR = [0.10, 0.15]). Furthermore, the factor of repeat/refresh interacted with age, indicating that older adults had a greater advantage of repeat over refreshed trials than young adults. Nevertheless, the repeat-refresh difference appeared for both young (Δ = 0.16, 95% HDR = [0.13, 0.18]) and older adults (Δ = 0.09, 95% HDR = [0.06, 0.12]).

The BGLMM revealed no credible evidence for a main effect of elaboration on WM performance, or for any of the interactions involving elaboration (see Table 3).

#### Long-term memory performance

The BGLMM revealed evidence for a main effect of repeat/refresh on LTM performance, but as with WM, there was an advantage for repeating over refreshing (see Table *4*). There was no evidence for any further interaction including the repeat/refresh factor. Hence, contrary to the findings of Johnson and colleagues, refreshing did not lead to better LTM than repeated reading. Note that the above pattern of results also holds for a lenient score of performance in the LTM task, counting all responses showing correct item memory (i.e. the target, same-list items, and other-list items) as correct responses.

Furthermore, the analysis of the LTM data revealed evidence for an interaction of elaboration with age (see Table *4*). Follow-up analyses of the interaction revealed that a beneficial effect of elaboration appeared only for young (Δ = 0.05, 95% HDR = [0.02, 0.06]), but not older adults (Δ = −0.01, 95% HDR = [−0.04, 0.3]). In sum, memory was better for trials with instructed elaboration than for those without, but only for the young and not the older adults. The above evidence speaks for an age-dependent beneficial effect of elaboration on LTM that is lost in older age.

To summarize, our results provide no evidence for an effect of refreshing on LTM for either age group; instead we replicated the benefit of elaboration on LTM but only for young adults.

### MVPA Results

#### Young adults

##### Repeat vs. Refresh vs. Elaborate vs. Refreshing with Elaboration

The classification scores for each individual were converted to a sensitivity score, accounting for both hits and false alarms, by computing the area under the ROC curve (AUC) for the four-way classification. For 24 of the 30 subjects classification of repeat, refresh, elaborate and refreshing-with-elaboration processes was successful (i.e., significantly better than chance with p < .05) in the predefined anatomical ROI (*MAUC* = 55.92 %, *SDAUC* = 3.48%, see Figure 3a). Data from the six remaining subjects were nevertheless included in the subsequent analyses, because excluding them would artificially restrict the range of classifier accuracy values that we used as predictors of behavior. The majority of the subjects (N = 19) showed classification in the whole mask of frontal–fusiform–parietal regions, and the remaining subjects (N = 5) showed good classification in the individual regions of the whole mask (Frontal: N = 3, Fusiform: N = 1, Parietal: N = 1). Notably, whole-brain classification was less successful than this ROI-based approached (M*AUC* = 54.79%, *SDAUC* = 2.69%, i.e., significantly better than chance with p < .05). The confusion matrix in Figure 3c depicts the frequencies for all classifications of each of the four conditions, including which misclassifications are more frequent than others. The matrix reveals that the conditions are overall well distinct, and if anything, the condition of refreshing with elaboration seems more similar to the visually equivalent refreshing condition.

**Figure 3.**
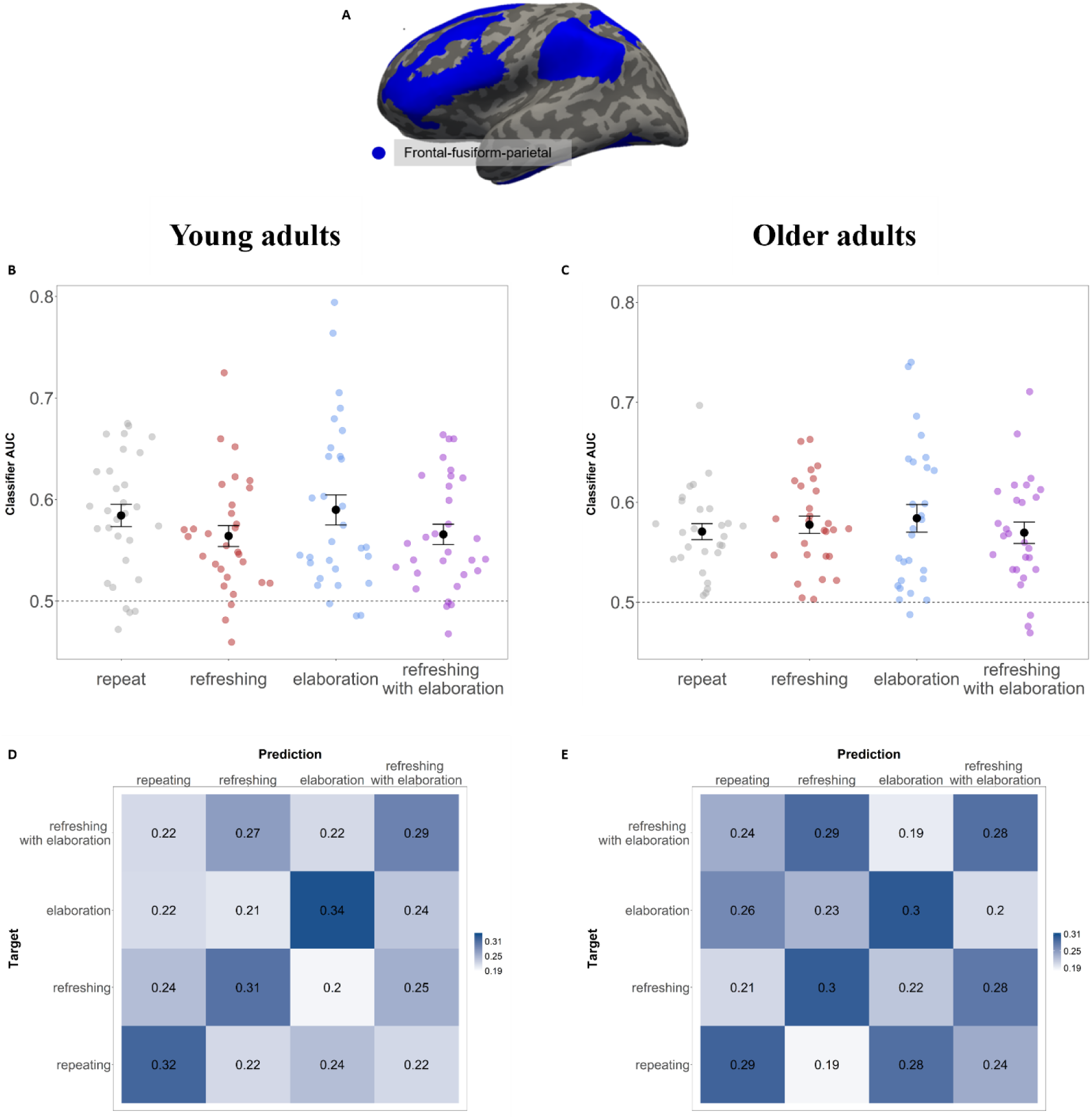
**A** Regions of interest for analysis include voxels in frontal cortex, fusiform gyrus, and parietal cortex (see Methods). **B and C** Classifier for Repeat vs. Refreshing vs. Elaboration vs. Refreshing-with-Elaboration in young and older adults, respectively. Decoding as indicated by the classifier AUC for the subjects. The error bars indicate the standard error. **D and E** Confusion matrix of the classifier accuracy for classifiers trained on Repeat vs. Refreshing vs. Elaboration vs. Refreshing-with-Elaboration in young and older adults, respectively. Chance performance is at 0.25.

##### Linking neural classification to memory performance

###### Refresh vs. Repeat

The classifier evidence values for refresh and repeat events were extracted from the four-way classifier and the resulting classification scores for each of the 30 individuals were converted to a sensitivity score, accounting for both hits and false alarms, by computing the area under the ROC curve (AUC) for the *refresh* vs. *repeat* classification. The mean classifier AUC for separating re-reading from refreshing was 0.557 (*SD* = .060, chance = 0.500). We performed a logistic regression relating the evidence values for the two processes on each trial to the WM outcome of that trial, and applied non-parametric bootstrapping using data sampled from all participants, in order to increase statistical power for this analysis (Efron, 1992; Lewis-Peacock et al., 2016) On each bootstrap iteration (n = 10,000), 30 participants were selected at random, with replacement, and their data were combined into a single *supersubject* for fixed-effects analysis. The stability of these effects across all iterations was analyzed to assess population-level reliability. For refreshing trials, WM outcome was operationalized in two ways: (1) as a binary outcome variable of whether the *refreshing processing benefit* (defined as the contrast between processed and not-processed triplet) of this trial was larger or smaller than the mean *repeat processing benefit* of this individual and (2) as a binary outcome variable of whether there was a *refreshing processing benefit*. None of the analyses revealed a significant relationship between WM outcome variables and neural separability (all ps > 0.31).

In summary, these results indicate that although repeating items benefited WM performance more than refreshing did, the neural pattern separability of these processes was not predictive of the size of this benefit.

###### Elaborate vs. Repeat

The classifier evidence values for elaborate and repeat events were extracted from the four-way classifier and the resulting classification scores for each of the 30 individuals were converted to an AUC sensitivity score for the elaborate vs. repeat classification. The mean classifier AUC for separating repeating from elaboration was 0.591 (SD = .090, chance = 0.500). To assess how the neural classification of the elaboration process relates to an individual’s task performance, we again used the repeat condition as a reference. We performed a logistic regression relating the evidence values for elaboration (relative to repeat) on each trial to the memory performance of that trial. Elaboration had no behavioral effect on WM, but instead showed a benefit for LTM. Therefore, our analysis focused on the behavioral contrasts in the LTM accuracy data: LTM outcomes were operationalized as a binary outcome variable of whether the *processing benefit of this elaboration* trial was larger or smaller than the mean *repeat processing benefit* of this individual. As shown in Figure 4a, stronger neural evidence for elaboration was related to a larger elaboration benefit in LTM (β = 1.12, p=.005). There was no such relationship for WM outcomes (β = 0.2, p=.449). The more elaborating on stimuli was neurally distinct from merely repeating those stimuli, the larger its beneficial effect on LTM beyond simply re-reading the words.

**Figure 4.**
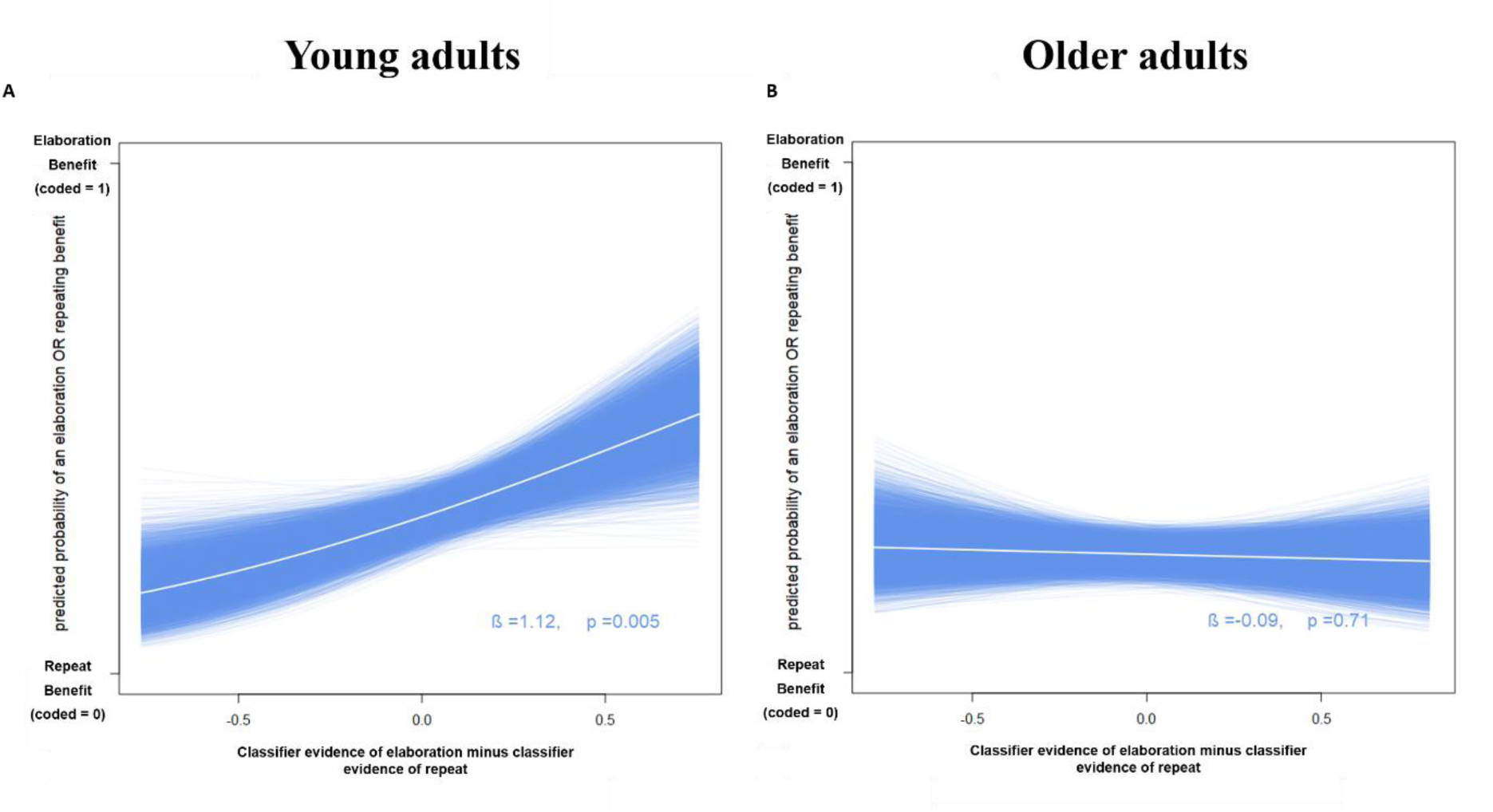
Neural evidence for elaboration predicts elaboration benefit in younger adults. On each of 10,000 bootstrap iterations, a logistic regression evaluated the classifier evidence for elaboration (relative to repeat) to the LTM elaboration benefit (relative to repeat) across each trial. The result from each bootstrap is visualized with a single blue line, and the mean is depicted in white.

#### Older Adults

##### Repeat vs. Refresh vs. Elaborate vs. Refreshing with Elaboration

The same analysis pipeline for the four-way problem in the young adults was subsequently applied to the independent sample of 27 older adults. For 17 subjects the classification of *repeat vs. refresh vs. elaborate vs. refreshing with elaboration* was significantly above chance in the in the predefined anatomical ROIs, with a mean classifier *AUC* of 0.5414 (*SD* =0.033, see Figure 3c). This result indicates that the four processes were neurally separable also in older adults, yet a smaller proportion of older subjects showed successful classification of the processes (17/27; 63%), compared to the young adult group (24/30; 80%). To gauge the strength of evidence for an age difference in classification success we carried out a Bayesian t-test comparing the AUCs of young adults to those of older adults; this test returned an ambiguous Bayes Factor of 1.25, meaning that there is neither evidence for nor against an age difference in the data.

Most of the older adults with above-chance classification (N = 13) showed good classification in the whole mask of frontal–fusiform–parietal regions, and the remaining subjects (N = 4) showed good classification in the individual regions of the whole mask (Frontal: N = 2, Fusiform: N = 2). Data from the ten subjects for which the cross-validation classification accuracy was not significantly above chance were nevertheless included in the subsequent analyses. As for the young adults, we also looked at the confusion matrix for the classification in older adults. Figure 3d depicts the frequencies for all classifications of each of the four conditions, including which misclassifications are more frequent than others. The matrix reveals that compared to the young adults, there were more misclassifications, such that the visually equivalent conditions of refreshing and refreshing with elaboration were frequently confused.

##### Linking neural classification to memory performance

###### Repeat vs. Refreshing

Equivalently to the analysis of the young adults’ data, the evidence values for refresh and repeat events were extracted from the four-way classifier, and the resulting classification scores for each of the 30 individuals were converted to a sensitivity score, accounting for both hits and false alarms, by computing the area under the ROC curve (AUC) for the refresh vs. repeat classification. The mean classifier AUC for separating re-reading from refreshing was 0.578 (SD = 0.062, chance = 50%). We performed a logistic regression relating the evidence values on each trial to the WM outcome of that trial and applied non-parametric bootstrapping using data sampled from all participants. As in the young adults, there was no significant relationship between either of the two WM outcomes (refresh benefit relative to repeat benefit, or refresh benefit alone) and the neural separability of the refreshing and repeating processes (both ps > 0.38).

In summary, these results replicate the findings in the young adults showing that although repeating items benefited WM performance more than refreshing did, the neural pattern separability of these processes did not predict the size of this benefit for the older adults.

###### Repeat vs. Elaborate

The mean classifier AUC for separating re-reading from elaborating was 0.57 (SD = 0.058, chance = 50%). We performed a logistic regression relating the evidence values on each trial to the WM and LTM outcome of that trial and applied non-parametric bootstrapping using data sampled from all participants. Just as in the young adults, elaboration had no behavioral effect on WM, but in contrast to the young adults, the older adults also showed no benefit of elaboration on LTM. Furthermore, an individual’s classifier evidence score for elaboration (relative to repeat) was unrelated to WM performance, measured by the *elaboration-minus-repeat benefit* (β = −0.04, p=.847). As shown in Figure 4b and in contrast to young adults, there was also no relationship to LTM performance, indicated by the same contrast (β = −0.09, p=.71).

## Discussion

The goal of the present study was to investigate to what extent elaboration and refreshing are separable processes, given prior reports of their neural overlap as well as their similar proposed roles for WM and LTM. We aimed at investigating whether refreshing and elaboration are distinct in their contribution to WM and LTM formation, whether they elicit separable neural activation patterns in fMRI, and how they relate to age-related memory deficits. We compared the neural and behavioral results of these processes to a control condition of re-reading (repeating) the words during the delay-period of a WM task. In the following, we discuss the effects of refreshing and elaboration on WM and LTM and we argue that these processes are distinct and have distinct consequences on memory performance in young and old adults.

### Are refreshing and elaboration distinct processes?

If refreshing and elaboration are two labels for the same process, then the pattern of behavioral effects should be the same for WM and on LTM, and the patterns of brain activity supporting these processes should be indistinguishable. In the present study, in a combined mask of a priori brain regions from frontal, temporal, and parietal lobes, we found successful differentiation of brain activity associated with repeated reading, refreshing and elaboration processes in a majority of the subjects. This neural evidence supports the assumption that refreshing and elaboration are implemented with distinct neural processes. In line with this differentiation, we also showed that the combined condition of refreshing with elaboration was discriminable from brain activation patterns in the other three conditions. Hence, combining two distinct processes resulted in a differentiable third neural state. As indicated by the AUC and further supported by visual inspection of the confusion matrix, the conditions of refreshing and elaboration were less frequently confused with each other than with the other conditions. This is additional evidence towards the distinctiveness of these processes. Further, as discussed in detail below, refreshing and elaboration resulted in distinct behavioral effects on tests of WM and LTM.

For a minority of subjects, the neural signatures were not classifiable significantly better than chance. There are three possible reasons for those subjects to be non-classifiable: First, this null result could mean that refreshing and elaboration are actually not different in these subjects. Second, it could be that these subjects were not classifiable in our ROI but recruited other brain regions for the processes instead. Finally, those subjects’ data could have included more noise, making it hard for the classifier to detect the signal.

Analysing the pattern of activity across voxels in the ROI including regions in frontal, parietal and temporal cortex is one way of assessing process-level differences in network activity. Future studies could extend our findings to further differentiate the neural implementation of working memory maintenance processes, for instance by measuring the functional connectivity patterns implemented during the different maintenance processes (Cohen et al., 2017).

### How does elaboration affect WM and LTM?

The elaboration process can be distinguished from mere re-reading by the accompanying distributed patterns of fMRI activity in task-relevant regions of the brain. Whereas elaboration showed no benefit for WM, it did facilitate LTM performance for young but not older adults. Accordingly, in young but not older adults, the degree of neural separability of repeating vs. elaboration was positively related to the individuals’ elaboration benefit in LTM: Greater separation between the neural processes of repeating and elaboration was associated with larger LTM benefits of elaboration within subjects (Figure 4). By contrast, the present results confirm prior studies showing that elaboration had no beneficial effect on WM. Our findings thereby fail to provide experimental support for the conclusion from previous studies which found that higher WM performance on complex-span tasks was correlated with individuals’ use of elaboration strategies such as imagery and sentence generation (Bailey, Dunlosky, & Hertzog, 2009; Bailey et al., 2008, 2011; Dunlosky & Kane, 2007). This discrepancy could be due to the present study using a simple-span paradigm and previous research relying on complex-span tasks. Alternatively, the correlation reported in previous studies might not reflect a causal effect of elaboration on memory – rather, participants who have good memory have more information in memory to elaborate on.

### How does refreshing affect WM and LTM?

We replicated the behavioral findings from Bartsch et al. (2018): Refreshed items were remembered better in a WM test than not-refreshed items, but repeating items benefited WM performance even more than refreshing.We further replicated the lack of a LTM benefit for refreshed compared to repeated items, which stands in contrast to previous fMRI studies (e.g. Raye, Johnson, Mitchell, Greene, & Johnson, 2005). One explanation for this could be the use of larger set sizes in the current study and in Bartsch et al (2018): Raye and colleagues asked their participants hold just one or two items in WM during refreshing, whereas in our studies, the subjects held six items in WM, refreshing three of them. Still, the processing benefit in WM indicates that the subjects were able to engage in a process that was beneficial to their memory performance. If this refreshing benefit arises through the build-up of LTM traces, we should have observed a refreshing benefit to LTM for the processed vs. non-processed words as well, but this was not the case. Therefore, we conclude that refreshing merely temporarily prioritizes the refreshed items in WM over the not-refreshed items.

### How do refreshing and elaboration contribute to age-related memory deficits?

A second goal of the present study was to investigate whether refreshing and elaboration and their impacts on memory are preserved in older adults. As in our young adult sample, the three processes of repeating, refreshing, and elaboration were neurally distinguishable in the predefined mask of frontal, parietal and fusiform regions for a majority of the older adults (Figure 3c). The comparison of these processes provided confirmatory evidence that, like young adults, older adults engaged these processes differently, but as in young adults, their degree of neural separability did not relate to subsequent WM performance.

There were hints in our data – albeit not backed by statistical support – that elaboration and refreshing were less distinctive in old than in young adults. The proportion of non-classifiable subjects was larger in the sample of older adults (10 of 27, 37%) than young adults (6 of 30, 20%). Further, the confusion matrix revealed more confusions between the conditions for older compared to young adults, especially regarding the *refreshing with elaboration* condition. This might indicate that the older subjects engaged similarly in those visually equivalent conditions, (a) either by ignoring the additional instruction to also form a vivid mental image at least in some trials of the *refreshing with elaboration* condition or (b) by also forming a mental image in the refreshing condition.

Assuming less distinctive maintenance processes in old adults would be in line with previous research suggesting a reduced distinctiveness of neural representations in older adults. Age has been associated with a decline in segregation of brain networks (e.g. Damoiseaux, 2017; Lorist, Geerligs, Renken, Saliasi, & Maurits, 2014; Morcom & Johnson, 2015), which further has been related to worse cognitive performance (Chan, Park, Savalia, Petersen, & Wig, 2014; Wang et al., 2010). In the context of the memory system, it has been shown that older adults’ representations in the visual cortex are less precise and this reduced precision was correlated with poorer subsequent memory performance (Zheng et al., 2017). Yet, a recent study (Carp, Gmeindl, & Reuter-Lorenz, 2010) suggests a more complex picture, showing reduced distinctiveness of visual cortical representations in older adults only at encoding– a condition similar to the repeat condition of the present study – whereas the distinctiveness of maintenance-related responses in prefrontal and parietal regions depended on the memory load. At low loads, older adults showed higher distinctiveness than younger adults, and at high loads, this pattern reversed, with higher distinctiveness in young adults. In the present study, some older adults might not have been classifiable due to their generally less precise neural representations contributing to more shared and imprecise activation patterns across the four conditions. The behavioral analyses revealed that, like young adults, older adults benefited from processing the items again in the refreshing condition, compared to the not-processed items, as shown in evidence against an interaction of age with the repeat/refresh by processing interaction (see Table 2). Using a similar paradigm to the present study, with cues directing refreshing to a subset of the memoranda after encoding, a recent study (Loaiza & Souza, 2019) confirmed that older adults are able to focus attention on no-longer perceptually available representations in WM, a critical component of refreshing, in conditions without distraction.

Similar to the young adults, refreshing had no benefit on LTM in older adults. This replicates the age-group specific findings of Johnson (2004), who also found no LTM benefit in older adults when comparing refreshing to re-reading. Refreshing was identified as an independent process, however, as it was neurally separable from both re-reading and elaboration. As refreshing was not related to LTM performance, even in the young adults, we conclude that deficits in refreshing are not responsible for the LTM deficit in older adults either. We directly replicated this behavioral finding in recent work (Bartsch & Oberauer, 2019).

The results on elaboration show that the fMRI classifiers were able to differentiate mere re-reading from elaborating in the older adults (Figure 3c&d). However, there was no LTM benefit of elaboration in older adults, whereas this effect was robust in the young group (Figure 2). We argue therefore that most of the older adults did perform some mental manipulation in the elaboration condition that was different from mere re-reading, but whatever it was did not affect their LTM performance. These results are in line with the *elaboration deficit hypothesis* (Smith, 1980): When they have to generate their own elaborations (here mental images), older adults do not benefit in the same way as young adults do. In our most recent work, we directly replicated this finding of an elaboration deficit in LTM of older adults (Bartsch & Oberauer, 2019).

Taken together, our results provide evidence that the LTM deficit of older adults might arise at least in part from a deficit in the process of elaboration. Future research should investigate whether age-related LTM deficits can be compensated for by *providing* richer representations – such as images or even sentences that contain or describe the to-be-remembered stimuli – rather than having the older adults *generate* the mental images, which they might find difficult to do.

## Conclusion

Our study revealed that the processes of repeated reading, refreshing, elaboration, and refreshing with elaboration are differentiable in brain activation patterns in both young and older adults. Elaboration can be neurally distinguished from mere reading. While it had no impact on WM, elaboration did improve episodic LTM for young adults, and the size of the benefit was related to the neural separability of elaboration: The more differentiated elaboration was from re-reading, the more elaboration benefited LTM. In contrast to the young adults, older adults’ episodic LTM did not benefit from elaboration, even though this process was neurally separable from reading. This suggests that older adults implemented a sub-optimal form of elaboration, and this may be a contributing factor to age-related deficits in LTM.

## Footnotes

1 The timing parameters where chosen based on a pilot experiment with young adults, which allowed participants to process the items in each of the 4 experimental conditions in a self-paced mode. The mean processing times (PT) where PT = 1419 ms in the repeat condition, PT = 1491 ms in the elaboration condition, PT = 1197 ms in the refreshing, and PT = 1198 ms in the refreshing with elaboration condition.

2 We chose this analysis because it tests the appropriate model for accuracy data. Analyzing accuracy data with a linear model (such as ANOVA) that incorrectly assumes normally distributed data is highly discouraged because it can lead to spurious results (e.g., Jaeger, 2008).

